# Odd-Even Oddball Task: Evaluating Event-Related Potentials (ERPs) during Word Discrimination Compared to Speech-Token and Tone Discrimination

**DOI:** 10.1101/2022.06.30.498354

**Authors:** Marcus Voola, An T. Nguyen, Welber Marinovic, Gunesh Rajan, Dayse Tavora-Vieira

## Abstract

The use of tonal and speech token auditory oddball tasks has been used abundantly in past literature; however, it has been argued that tasks using non-word sounds fail to capture the higher-level ability to interpret and discriminate stimuli based on meaning which are critical to language comprehension. As such this study aims to examine how neural signals associated with discrimination and evaluation-processes (P3b) from semantic stimuli compare with those elicited by pure tones and speech tokens.

This study comprises of two experiments, both containing thirteen adults (Exp 1: (Mean(SD)_age_ = 25.20(3.89) years and Exp 2: Mean(SD)_age_ = 25.3(3.79) years) with normal hearing in both ears (PTA ≤ 20dB HL). Scalp electroencephalography and auditory event related potentials was recorded in free field whilst they completed three different oddball tasks: 1) Tones, 2) Speech tokens and 3) Odd/Even numbers. In experiment two, the duration of each stimulus was the same.

P3b peak latency was significantly different between all three tasks, in both experiments. P3b amplitude was identified to be sensitive to reaction time, with tasks that have a large reaction time variability resulting in the P3b amplitude to be smeared out, thereby reducing the amplitude size.

The findings from this study highlight the need to take into consideration all factors of the task before attributing any effects to any additional process such as semantic processing and mental effort. Furthermore, it highlights the need for more cautious interpretation of P3b results in auditory oddball tasks.

## Introduction

In 1980 Kutas and Hillyard identified a neural correlated of language processing using Event Related Potentials (ERPs). Even Related Potentials capture the cortical responses after the presentation of auditory stimuli, which can be in the form of tones, speech tokens or words (Balkenhol et al., 2020). Recording the brain activity in response to auditory stimuli provides a method for understanding how the brain process various auditory inputs and what factors are important for the processing of sound. The auditory ERP, P3b is thought to quantify an individual’s ability to discriminate and interpretate auditory stimuli (Polich, 2007).

The P3b is characterised by parietally distributed positivity occurring at a latency range between 300 ms to 600 ms after stimulus presentation. One method to elicit the P3b is by using an oddball paradigm which consist of two types of stimuli, an infrequent (target) stimulus and a frequent (standard) stimulus, whereby the target stimuli are characterized by a unique feature (Polich, 2007). The P3b positivity is enhanced upon the presentation of the target stimuli in the oddball paradigm when compared with the presentation of the standard stimuli (Didoné et al., 2016). The P3b has been suggested to reflect the decision-making ability of an individual and subsequently activation of stimulus response links. If this decision-making hypothesis is true, then it is expected that P3b amplitude larger and latency will be delayed with more complex tasks as the decision now takes demands greater higher order processing capabilities (Polich, 2007; Kelly and O’Connell, 2013). This rationale is further supported by the link between P3b and decision making is supported by the alignment of the P3b deflection to the onset of the response (Sassenhagen et al., 2014).

Traditionally, studies using P3b have looked at discrimination of tones, but more recent studies use more complex word-like stimuli, arguing that the latter is more naturalistic and contains semantic aspect which is missing from tones and may be better for language research (Verleger et al., 2014; Li et al., 2019). Speech tokens have also been used to investigate the higher order processing capabilities of an individual. Similar to the establishment of tonal stimuli, oddball paradigms incorporating speech tokens have also been adapted to be more complex by using speech tokens that are more phonetically similar. The P3b of speech tokens that were more phonetically similar (ie /ba/ vs /ga/ is more phonetically similar than /ba/ vs /di/) had delayed P3b latency and large amplitude. These finding highlights that delayed and enhanced (more positive) P3b may suggest that sounds which are phonetically similar require more time and cognitive resources to discriminate (Didoné et al., 2016).

Whilst tonal and speech-token tasks have been used abundantly in the past, they fail to capture the ability to interpret and discriminate stimuli based on meaning. This is attributed to both tasks involving the discrimination of the physical characteristics of the sound (e.g., frequency and fast-changing temporal qualities associated with articulation). As such any deficits in the processing of auditory stimuli may not be revealed by tonal or speech token oddball tasks that only require subjects to differentiated based on the physical properties and not the meaning of the sound.

Words have been used in oddball paradigms to elicit these ERPs. In 2016, Finke et al. used an oddball paradigm that required subjects to semantically classify words as either living or non-living entities(Finke et al., 2016). Greater stimulus complexity, in the form of differentiating stimuli based on meaning rather than just physical properties, may result in more complex neural circuity being engaged. These additional circuits include retrieving word meanings from our mental lexicon and the circuits involved in categorising words based on these meanings. This rationale is supported by past literature which identified that the more cognitive needed to classify words in an oddball paradigm results in delayed P3b latency and larger amplitude, when compared with tonal oddball tasks (Kotchoubey and Lang, 2001). This finding suggests that the use of a semantic oddball task requires additional processing to categorise the stimuli.

### Current Study

This study comprises of two experiments, both of which share the same aim. Both experiments aim to develop a semantic oddball task which requires participants to discriminate sounds based on their meaning rather than just the physical properties of the sound. In addition, both experiments aim to examine how neural signals associated with discrimination and evaluation-processes (P3b) from semantic stimuli compare with those elicited by pure tones and speech-like sounds (speech-tokens). We will be comparing the size of the semantic evoked ERPs with simpler and already established auditory oddball paradigms, tonal and speech tokens.

It is hypothesized that the additional processing required to discriminate between the semantic odd/even tasks will result in a larger and delayed P3b when compared with the tonal and speech token task. Upon analysis of the first experiments results we identified that whilst there was a difference between the three tasks, these differences could be attributed to the lack of control for critical latency (the earliest point after stimulus-onset when the target stimuli can be discriminated from standard stimuli) and stimulus duration rather than a semantic component. Experiment 2 was developed to validate these findings.

## Materials and Methods

This study consisted of two separate experiments. In both experiments, each participant completed three versions of the acoustic oddball task (tonal, speech-token and odd-even numbers). In experiment 2, stimulus durations were adjusted to correct for differences across the three tasks. While this correction modified ERP latencies, it did not completely account for task effects observed in experiment 1 (See Discussion). In the following sections, both experiments will be described together, and distinctions will be highlighted in the relevant sections.

### Participants

Participants were healthy adult volunteers recruited from the Audiology department at Fiona Stanley Hospital. Experiment 1 consisted of thirteen participants (Mean(SD)_age_ = 25.20(3.89) years, 6 females, 7 males). Experiment 2 also consisted of thirteen participants (Mean(SD)_age_ = 25.3(3.79) years, 5 females, 8 males). Four participants completed both experiments 1 and 2. All participants had their hearing thresholds tested and were all within the normal hearing range (less than 20 dB HL in all frequency ranges from 0.5 Hz to 8 kHz in both ears). The mean (SD) unaided air pure tone average (PTA) from 0.5 Hz to 8 kHz in experiment 1 was 4.95(3.60) dB HL in the left ear and 4.61(2.92) dB HL in the right ear. In experiment 2 the PTA was 6.15(2.41) dB HL in the left ear and 5.29(2.13) dB HL in the right ear. All participants provided written-informed consent before the beginning of the experiment. Ethical approval for this study was obtained from the South Metropolitan Health Ethics Committee (Approval Code: 335).

### Procedure

For both expeirments, participants were seated in a dimly lit sound attenuated room, facing a wall. Participants were fitted in an EEG cap and were provided with verbal instructions before completing three experimental tasks (see Experimental Tasks section). Self-paced breaks were taken between tasks, and task order was counter-balanced across participants. The entire experiment took around 30 minutes. A schematic diagram of the experimental set up is depicted in figure 1.

**Figure 1.**
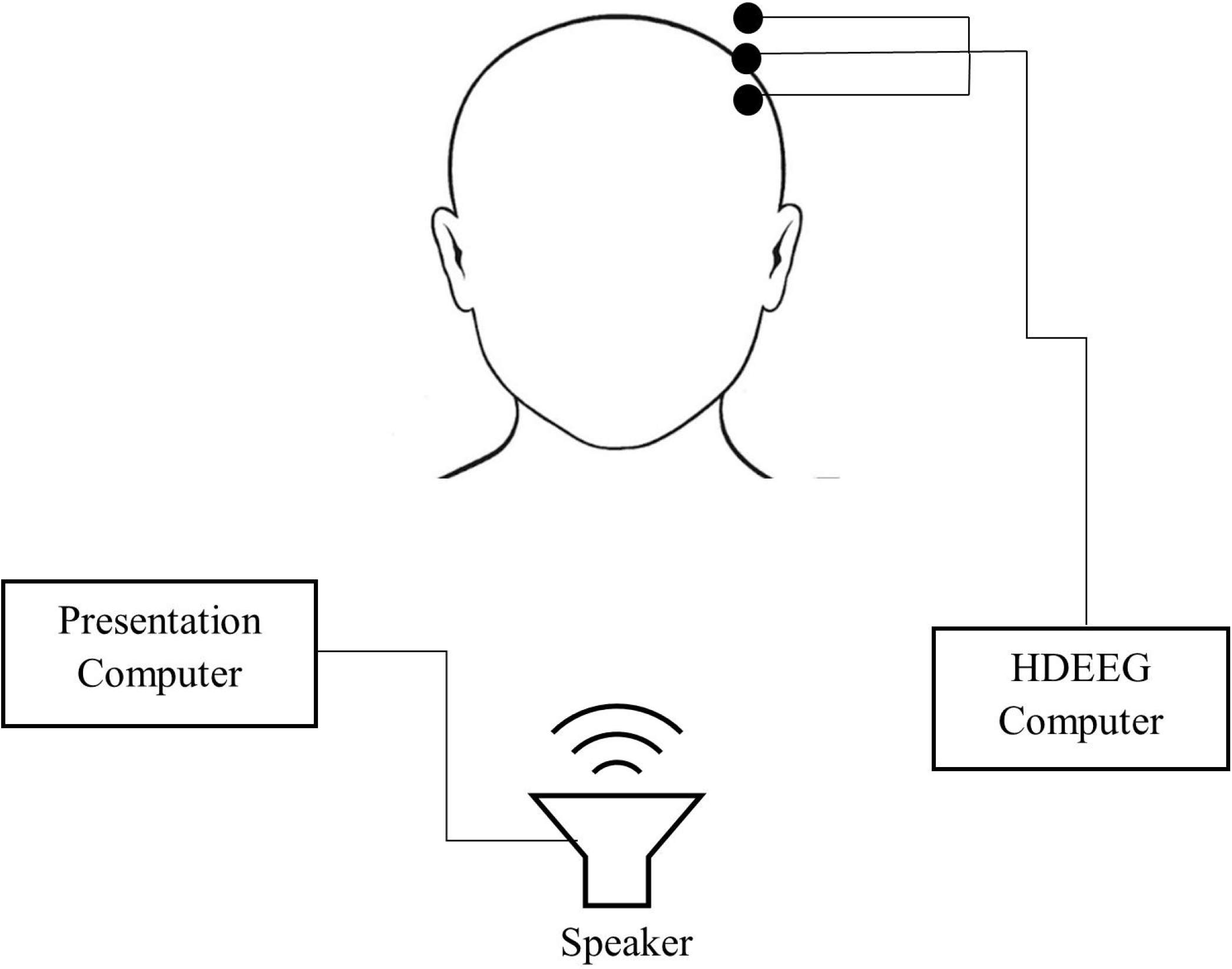
Schematic diagram of the set up used in both experiment 1 and 2. Testing was conducted in a sound attenuated booth.

### Experimental Tasks

Each experiment consisted of three acoustic oddball tasks with different stimuli (pure tones, speech-tokens and odd-even numbers). In each task, participants completed 240 trials where they were presented with an acoustic ‘standard’ or ‘target’ stimulus, and their task is to respond as soon as they hear the target stimulus by pressing a button on the response-pad with their right-thumb. Standard and target stimuli were presented on 80% (192) and 20% (48) of trials, respectively, and trials were presented pseudo-randomly such that a target stimulus would not be presented on two consecutive trials. All stimuli were presented in free-field at a calibrated intensity of 55-60 dB SPL using EDIFIER M1250 Multimedia Speakers, with an inter-stimulus interval of 1800 ms. Cogent 2000 and Psychtoolbox-3 functions in MATLAB were used to control the experiment. Reaction time is the time of button press relative to stimulus onset The pure tone and speech-token tasks each consisted of two stimuli, and the odd-even numbers task consisted of eight stimuli. In the pure tone task, 1 kHz and 2 kHz pure tones were used. In the speech-token task, synthesised /ba/ and /da/ sounds were used. These speech-tokens were obtained from National Acoustic Labarotaries (NAL) and were first used in a 2008 study conducted by Johnson et al.(Johnson et al., 2008). In the odd-even numbers task, eight pre-recorded numbers from one to nine were used. The number seven was omitted because it was the only number to contain two syllables. These recordings were obtained from NAL and were recorded by a mature female Australian speaker. These speech files were recorded with the purpose to be used in a telephone-based speech-in-noise test called ‘Telescreen’(Dillon et al., 2016). These recordings were modified in Audacity to reduce the discrepancy in stimulus duration between numbers while maintaining intelligiblity. In experiment 1, the duration of pure tones were 100 ms, the duration of speech-tokens 170 ms, and the average duration of odd-even numbers was 400 ms. In experiment 2, the duration of pure tones and speech-tokens were lengthened using Audacity, to match average duration odd and even numbers.

### Acquisition and Pre-Processing of Electrophysiological Data

Electrophysiological data were continuously recorded for the duration of each task. Data were acquired using the Micromed™ SD LTM EXPRESS system, a SpesMedica cap (SpesMedica™, Genova, Italy) and Gilat Medical ERP software (Gilat Medical Research & Equipment Ltd, Karkur, Israel), at a sampling rate of 1024 Hz with an online low pass-filter of 40 Hz. Data was recorded from 59 Ag/AgCl scalp electrodes arranged according to the 10-20 system with additional electrodes placed under the right infraorbital region to monitor eye-movement, a reference electrode placed on the middle of the chin, and a ground electrode placed on the right mastoid. All electrode impedance was kept below 5 kΩ for the duration of the recording.

MATLAB 2020a was used to process the data. A semi-automated procedure was used consisting of functions from the plug-ins EEGLAB (Delorme and Makeig, 2004), PREP pipeline (Bigdely-Shamlo et al., 2015), clean_rawdata() plugin, AMICA (Palmer et al., 2011) and ICLabel plugin (Pion-Tonachini et al., 2019). The removeTrend() from the PREP pipeline plugin was used to linearly detrend the data using a high-pass 1 Hz FIR filter (step size = 0.02). The cleanLineNoise() from PREP pipeline plugin was used to remove 50 Hz line noise and harmonics up to 500 Hz. The pop_clean_rawdata() was used to determine noisy channels. The pop_interp was used to interpolate noisy channels spherically. EEG data was then down sampled to 250 Hz. The data was demeaned, and a 30 Hz low-pass filter (order = 100) was applied using the pop_eegfiltnew(). The clean_asr() was used to correct for artefacts using the artefact subspace reconstruction method (SD = 100).

Initially, the data were epoched from -200 to 1000 ms relative to stimulus-onset. However, further investigation highlighted that task-effects may be driven by difference in ‘critical point’, the earliest point after stimulus-onset when the target stimuli can be discriminated from standard stimuli. For tones, the critical point was 0 ms (i.e., at sitmulus onset). For speech tokens, the critical point was 40 ms. For odd-even numbers, the critical point ranged from 30 to 250 ms. The critical point determined manually by two normal hearing individuals. As such, we also extracted -400 to 1000 ms epochs time-locked to the critical point of each stimulus. Independent component analysis (ICA) of the data was conducted using AMICA (2000 iterations) on down sampled data to 100 Hz (Palmer et al., 2011). The number of independent components extracted were adjusted for the data rank. ICLabel functions were used to classify and remove independent components that were eye movement, muscle, heart, line noise, or channel noise with > 70 % confidence. The data was baseline correct was to the pre-stimulus interval. Trials with activity exceeding 100 μV were flagged for exclusion for further analysis.

### Measurment of Event-Related Potentials

We measured the mean amplitude of P3b at the single trial level, on waveforms time-locked to stimulus-onset and the critical-point. P3b amplitude was measured at the centroparietal mid-line electrode (CPz) where the P3b was most prominent. P3b was also measured over a 40 ms interval around the positive peak of the grand average waveform between 300-600 ms.

Peak latency of the P3b were measured at the level of the participants grand average waveform, time-locked to both stimulus-onset and the critical-point. Measurment at the level of the participant was undertook after single trial level measurments resulted in large variability for peak latency. P3b peak latency was measured as the time-point of maximum amplitude between 300-800 ms at CPz.

### Reaction Time

Reaction time was the time taken from target stimulus onset for the subject to respond to the presentation of the target stimuli. To account for differences in critical latency between the three oddball tasks, adjusted reaction time was cacluated at a trial level by subtracting the critical latency for the presented target stimuli from the subjects reaction time.

### Stimulus and Response-Alligned ERP image

To illustrate the relation between P3b, stimulus-onset and reaction time, we generated ERP image plots (See Fig. 5B) depicting singletrial EEG activity focusing on the P3b effect (Target-Standard difference waveforms at CPz). The EEG data plotted represents a pool of trials from all participants, which was sorted by reaction time and plotted as a colormap (blue = negative, red = positive). Trial level difference waveforms were achieved by subtracting the grand-average standard ERP from each target trial waveform. We also indicated the time of stimulus and response-onset by overlaying black solid and red dashed lines, respectively.

**Figure 2.**
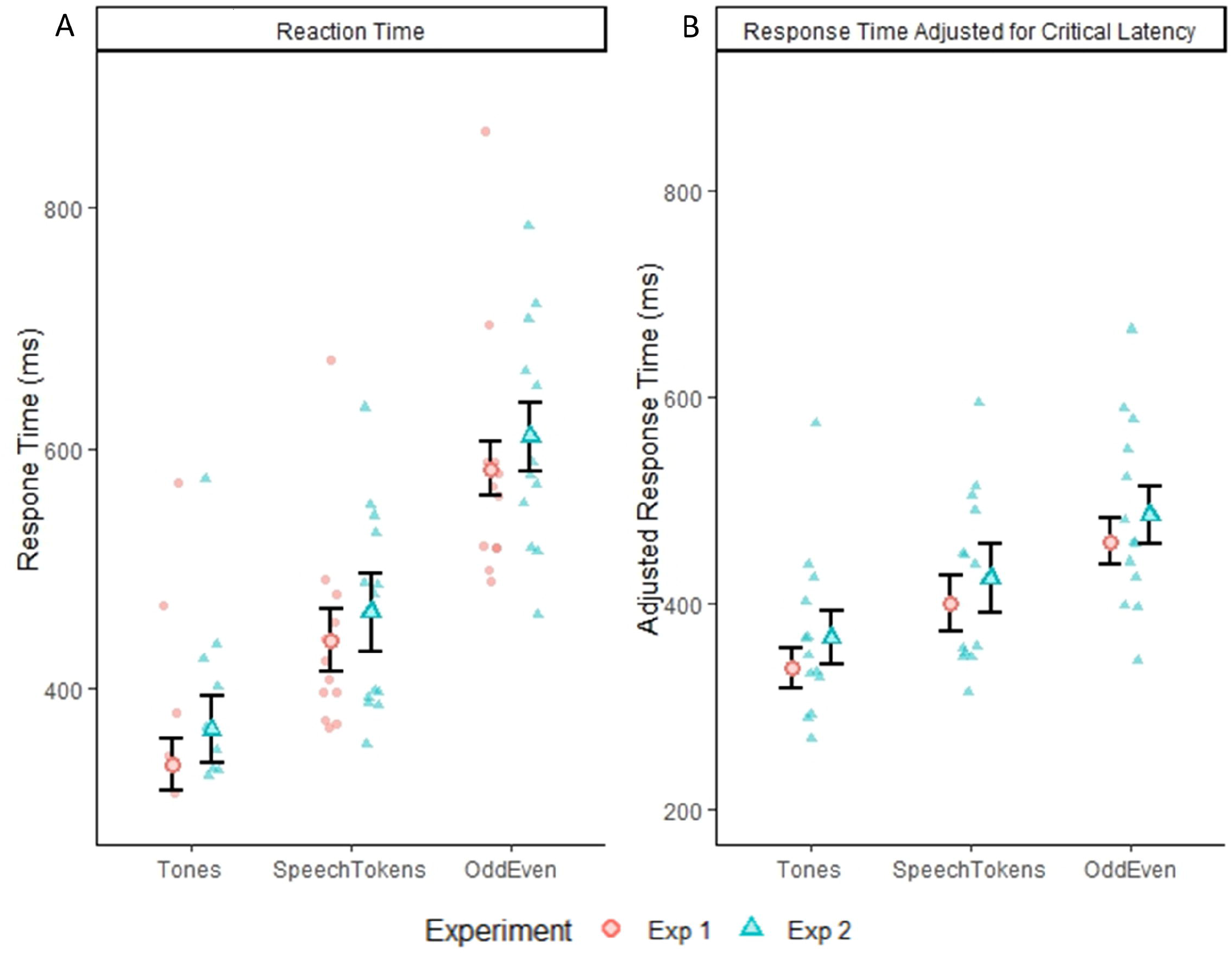
Panel (A) depicts the group average response time with within subject error bars for all three tasks from experiment 1 and 2. Panel (B) depicts the group average response time adjusted for critical latency. Error bars reflect standard error of the mean

**Figure 3.**
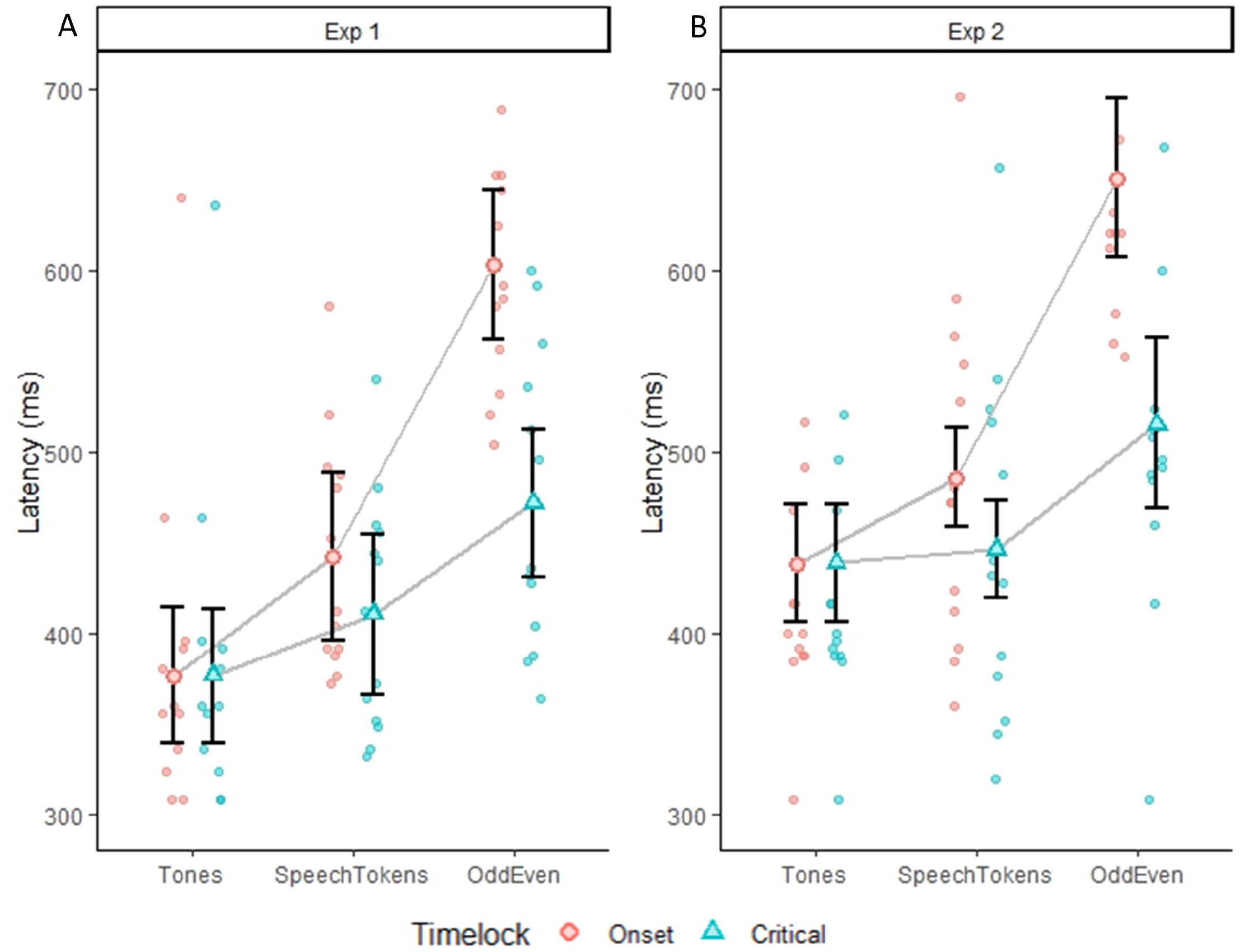
Plots depicting the average P3b peak latency for all three tasks average on a subject level. Error bars represent the standard error of the mean. Panel (A) depicts the difference in P3b peak latency for experiment 1 when time locked to onset (red dots) and critical latency (blue dots). Panel (B) depicts the difference in P3b peak latency for experiment 2 when time locked to onset (red dots) and critical latency (blue dots). Faded red and blue dots represent each subject P3b latency.

**Figure 4.**
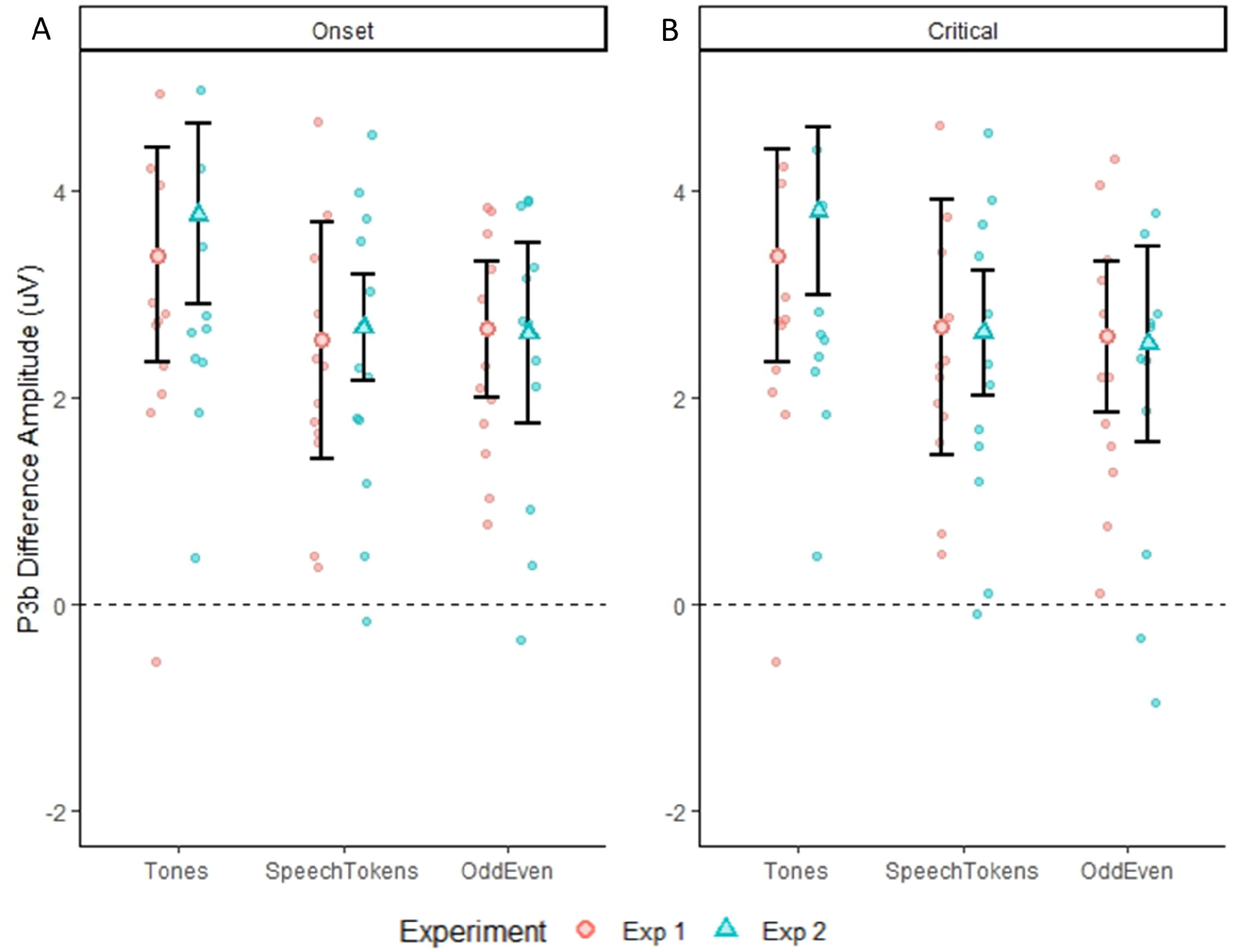
Differences in P3b amplitude for both experiments for all three tasks calculated at a single trial level. Error bars represent standard error of the mean. Panel (A) shows the difference in the three tasks for both experiments when time locked to onset latency. Panel (B) shows the difference in the three tasks for both experiments when time locked to critical latency.

**Figure 5.**
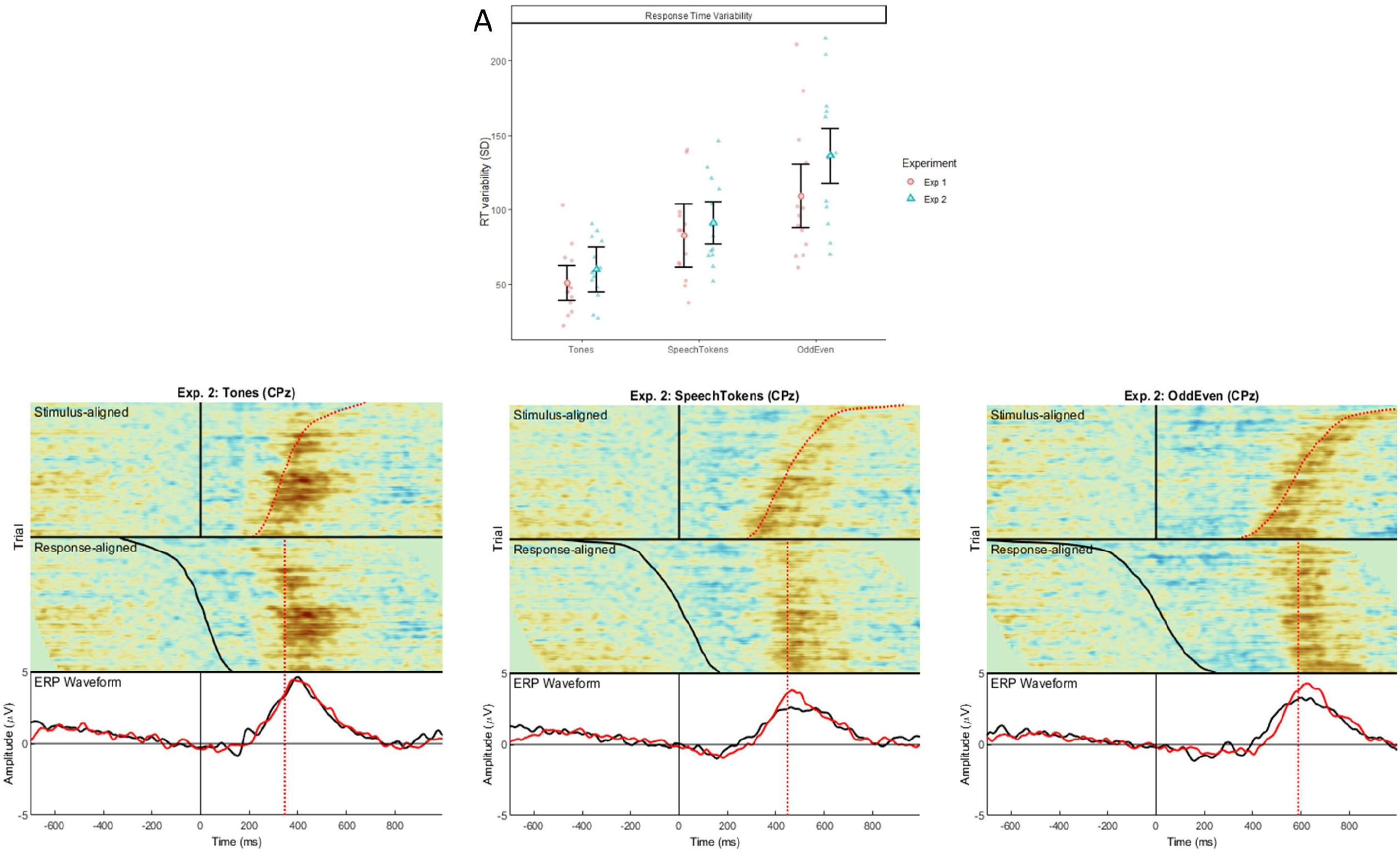
Panel (A) illustrates the difference in response time variability for each three tasks both experiments 1 and 2, calculated at a single trial level. Error bars represent standard error of the mean. Shaded red circles and blue triangles represent the average reaction time variability for each subject. Panel (B, C and D) shows the stimulus aligned waveform (i.e., time locked to stimulus onset) and response aligned (i.e., time locked to the subject’s reaction time) for tones, speech tokens and Odd/Even respectively. The black ERP waveform represents the waveform generated from the stimulus aligned data and the red ERP waveform represents the response aligned data.

In the ERP image, we represented the data in two ways; ‘stimulus-locked’ and ‘response-locked’. For stimulus-locked, the ERP data was aligned to stimulus-onset (illustrated by the vertical black line at 0 ms on the top-half of the image) and it can be seen that RT varies across trials (illustrated by the curved red line ∼ 400 ms). Most notably, it can be seen that the positivity in the EEG signal (shown by the red of colormap) roughly coincides with RT. In other words, the P3b activity seems to show a response-alignment. To further highlight this point, we used ‘response-locked’ plots which re-aligns the same EEG data to time of response-onset (illustrated by the vertical lines between 350-600 ms on the bottom-half of the image). In this plot, it can be seen that positivity is no longer ‘smeared’ over time and is clustered around the red dashed line, illustrating that the P3b shows a response-alignment.

Under the EPR image are two grand-averaged waveforms computed from the stimulus and response-locked ERP data. For Speech-Token and Odd/Even tasks, it can be seen that the amplitude of stimulus-locked data is smaller than response-locked data because the timing of the P3b is more varied in the stimulus-locked data. This illustrates that differences in P3b amplitude are a by product of differences in RT variability (See Fig. 5A) as opposed to a true difference in amplitude across tasks.

### Statistical Analysis

All statistical analysis was conducted using R statistics and R software. A linear mixed model analysis was employed using “lmer” function from the “lmerTest” package. For the N2N4 analysis we modeled the amplitude with trial type, task and the interactions has fixed effects, with participant intercepts modeled as a random effect. For P3b we moduled amplitude using the same method. The results of the model were presented as F values using the the Satterthwaite’s approximation method with the ‘ANOVA’ function. We used “r2beta” to calculate the effect size. Main effects and interactions were evaluated by comparing the estimated marginal means using the “emmeans” function from the “emmeans” package. The result of these pairwise categorical comparsions were presented as t-ratios (mean difference estimate divided by standard error) with degrees-of-freedom estimated using the Kenward-Roger method.

## Results

### Overview

In this section, the results obtained from both experiment 1 and 2 are presented. The P3b peak amplitude and latency figures are presented to illustrate the difference observed between the three different oddball tasks. These illustrations are repeated twice for each experiment, once looking at the effects when time locked to onset latency and second looking at the effects when time locked to critical latency. In addition, differences in reaction time between the three tasks for both experiments are also depicted. Overall, the results from both experiments suggest that the difference between the Odd/Even oddball task when compared to the speech token and tonal oddball task can be attributed to either physical differences in the stimuli (critical latency and stimulus duration) or a semantic component.

### Reaction Time

The reaction time for all three oddball tasks across both experiments is displayed in figure 2A. Pairwise comparisons identified a significant difference for comparisons between all three tasks: tones vs speech tokens (t(3692) = -24.164, *p* = <0.0001), speech tokens vs Odd/Even (t(3692) = -34.669, *p* = <0.0001) and tones vs Odd/Even (t(3692) = -58.860, *p* = <0.0001). Reaction time was also calculated considering the difference in critical point between conditions (Fig. 2B). Despite accounting for difference in critical latency, pairwise comparison of all three oddball tasks revealed a significant difference between all three tasks: tones vs speech tokens (t(3692) = -13.892, *p* = <0.0001), speech tokens vs Odd/Even (t(3692) = -13.928, *p* = <0.0001) and tones vs Odd/Even (t(3692) = -27.826, *p* = <0.0001).

### P3b Latency

In experiment 1, P3b peak latency when time locked to onset latency identified that in all three oddball tasks had significantly different P3b peak latency (t(60) = 37.341, *p* = < 0.0001) (Fig. 3A): tones vs speech tokens (t(24) = -2.222, *p* = 0.0360), speech tokens vs Odd/Even (t(24) = -5.445, *p* = < 0.0001) and tones vs Odd/Even (t(24) = -7.666, *p* = <0.0001). When time locked to critical latency, a significantly shorter P3b peak latency was only observed for tones when compared with Odd/Even ((t(24) = -3.286, *p* = 0.0094).

When stimulus duration was controlled for in experiment 2, P3b peak latency was still when time locked to onset latency was significantly delayed for Odd/Even when compared with both tones (t(24) = -8.753, *p* = <0.0001) and speech tokens (t(24) = -6.799, *p* = <0.0001) (Fig. 3B). When time-locked to critical latency, similar effects were observed except the magnitude of the difference was reduced. P3b peak latency for Odd/Even was still significantly delayed when compared with tones (t(24) = -7.666, *p* = 0.0173) and speech tokens (t(24) = -2.727, *p* = 0.0235).

### P3b Amplitude

P3b amplitude time-locked to stimulus onset and averaged over both experiments elicited a significantly larger amplitude for tones when compared with speech tokens (t(3477) = 4.934, *p* = <0.0001) and Odd/Even (t(3476) = 4.782, *p* = <0.0001). Likewise, P3b amplitude time-locked to critical latency indicated a significantly larger amplitude for tones in comparison with speech tokens (t(3445) = 4.700, *p* = <0.0001) and Odd/Even (t(3444) = 5.334, *p* = <0.0001). Figure 4A and 4B illustrate the P3b waveform morphology of both experiments and time-locks.

### Response Variability and P3b Response-Alignment

Reaction time variability was also calculated to examine the response-alignment of the P3b signals across the three tasks and both experiments (Fig. 5A). Critical latency of target trials was subtracted from the subject’s response time at a trial level. Reaction time variability was significantly smaller for tones when compared with speech tokens (t(48) = -3.720, *p* = 0.0005) and Odd/Even (t(48) = -8.485, *p* = <0.0001). Speech tokens was identified to be significantly smaller than Odd/Even (t(48) = -4.765, *p* = <0.0001).

Comparison of stimulus and response aligned ERPs are illustrated in figure 5B, 5C and 5D for tones, speech tokens and Odd/Even respectively. Response aligned ERPs indicate that the P3b amplitude for speech tokens and Odd/Even is suppressed due to the high reaction time variability. On the other hand, the small reaction time variability in the tonal task elicits a larger P3b due more consistent response from trial to trial. We analysed the effect of task on the stimulus and response aligned P3b amplitude data (Fig. 6) and identified that the significant effect of task in the stimulus aligned data (*F*(2,1691.4) = 4.05, *p* = 0.018*) is no longer present when you align the response data (*F*(2,1691.2) = 0.40, *p* = 0.668).

**Figure 6.**
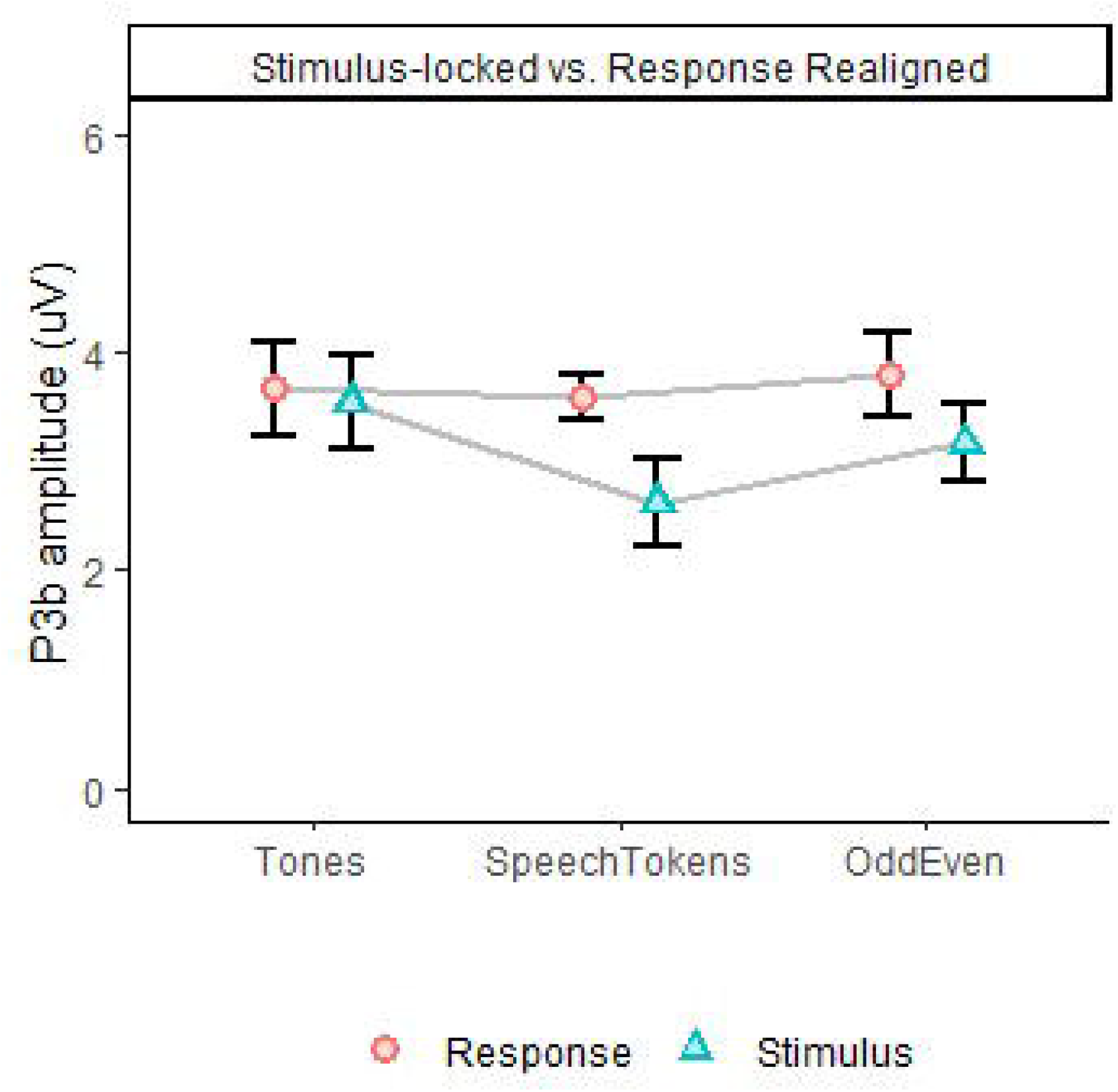
P3b amplitude for all three tasks in Experiment 2, depicting differences in task effect on stimulus and response aligned data. Error bars represent standard error of the mean.

## Discussion

Recently, it has been argued that tonal oddball tasks lack the semantic component which are critical to language processing, therefore making it a bit far-fetched from real world application. In addition, there are claims that semantic oddball tasks using words may engage additional semantic processing which might enhance the associations between ERP responses and measures of language and comprehension. Furthermore, this may lead to more specific treatment outcomes in patients with hearing difficulty. To address these claims, this study had two aims. Firstly, we aimed to develop a semantic oddball task which force participants to discriminate auditory stimuli on their meaning rather than just differences in the physical properties of the sound. Secondly, we aimed to compare this task against more established oddball paradigms that use tonal and speech token stimuli. We hypothesized that with increasing stimulus complexity, response time would increase and P3b latency and amplitude would be delayed and enhanced respectively. All of which indicating increased mental effort associated with evaluating increasing complex stimuli.

In experiment 1, we identified an increase in reaction time and P3b latencies for increased stimulus complexity (tones < speech tokens < odd/even). Amplitude of P3b, did not follow the same trend but rather it was identified to be larger for simple stimuli (tones) when compared with complex stimuli (speech tokens and odd/even numbers). In trying to understand why this result was observed, we identified that the difference in critical latency and stimulus duration between the three oddball tasks could be an explanation.

The critical latency of all three tasks were not the same, with tones having a critical latency of 0 ms, speech tokens having a critical latency of 40 ms and odd/even having a critical latency ranging from 60 to 250 ms. For example, differentiating between pure tones can be done by the subject as soon as they hear the stimulus, as it only requires them to detect the difference in frequency. Similarly, speech tokens can also be differentiated in the same manner as tonal stimuli as the subjects are differentiated the sounds when they are able to hear the /d/ or /b/ phoneme. On the other hand, differentiating between odd and even numbers will take longer even if the subject has detected the sound. This is highlighted in a subject trying to differentiate between the numbers ‘four’ and ‘five’. Both numbers begin with a /f/ phoneme, so even though a subject detects this sound they still must wait for more auditory information before they can differentiate the two numbers. To account for these differences, the data from experiment 1 was re-analysed time-locked to the critical latency. When aligned to critical latency there was no significant impact on the reaction time, P3b latency and P3b amplitude with the same trend in results being observed. The size of the difference was reduced but the effect was still significant.

The second possible explanation for the trend in results was the difference in stimulus duration across all three oddball tasks. This rationale is supported by previous studies who have identified that endogenous potentials such as ERPs, are affected by stimulus properties such as intensity, pitch and stimulus duration (Alain and Winkler, 2012). This led to the development of experiment 2, which contained the same three oddball tasks as experiment 1 but the stimulus duration of the tonal and speech token oddball task was matched to the duration of the odd/even task. Despite the control of stimulus duration, we identified that the same trend in data from experiment 1 was identified. Reaction time, P3b peak latency and P3b amplitude were all significantly different, with the effect size being reduced when data was analysed time locked to critical latency but still being significant. The two explanations, critical latency and stimulus duration, were both found to have no significant effect on the results. This indicates that the results observed in the present study cannot be attributed to differences in critical latency and stimulus duration.

The enhancement of P3b amplitude for tones in comparison with speech tokens and odd/even was surprising. Our initial hypothesis thought that the P3b amplitude would be enhanced for increasing stimulus complexity, however we observed the opposite. Further analysis indicated that the large variability in reaction time could be an explanation for the P3b amplitude findings. As such, reaction time variability was calculated which identified that the variability was the smallest in tones, then speech tokens and then largest odd/even numbers. To account for these differences, data from experiment 2 was re-analysed time-locked to the response time (Fig. 5A). We found that once accounting for variability in response time, the P3b amplitude of both speech tokens and odd/even numbers increased. The P3b amplitude for tones stayed the same as there was very little difference in reaction time variability. This finding is in line with past studies which have identified that variability in reaction time has a greater effect on P3b amplitude than P3b peak latency (Ramchurn et al., 2014; Verleger, 2020). Ramchurn et al (2014) identified that within the same subject recording, response times that were faster elicited a significantly larger P3b amplitude than slower response times (Ramchurn et al., 2014). Thereby the large variation in reaction time seen in the odd/even task results in the effect of P3b amplitude to be smeared out rather than a concentrated peak being produced. This finding is supported by the significant differences in peak waveform morphology between the three tasks shown in figure 5. Follow-up analysis identified that the effect tasks is no longer significant when aligned to the response (Fig. 6).

The variability seen in the reaction time for the odd/even task could be attributed to the differences in the number of stimuli per task. More specifically, in the odd/even task there are four different targets and standards, each having a unique critical latency, whereas in the tones and speech token task there is only one standard and target for each task. The greater variability in critical latency for the odd/even task is a possible explanation for the variability in reaction time. Other factors such as the spectral quality of the sound and the semantic complexity cannot be ruled out but can be thought to contribute to the variability. Future studies should look to control for the number of stimuli per task when comparing stimulus complexity.

Taken all together, this study demonstrates that increasing stimulus complexity results in an increase in reaction time, reaction time variability and P3b latency. We identified that critical point and stimulus duration can only partly explain the effect of reaction time and P3b latency. Other factors such as differences in the spectral qualities of stimuli, semantic complexity, and the difference in the number of stimuli per task could also have contributed to the results. Additionally, variance in reaction time was identified to explain the P3b amplitude findings. Despite difference in stimuli, the morphology of the ERP waveforms was similar between tasks – suggesting that similar mechanisms are used between the tasks.

The findings from this study highlights the importance of considering each aspect of the task before attributing effects to additional processes such as semantic processing and mental effort – especially for amplitude. The study calls for more cautious interpretation of P3b results in semantic oddball tasks compared to simpler tasks. Future studies should look to examine the effect that the number of stimuli per task has on reaction time variability and P3b latency

## Funding Sources

This research was partially supported by the Australian Government through the Australian Research Council’s Discovery Projects funding scheme (project DP180100394) awarded to WM. This project did not receive any other specific grant from funding agencies in the public, commercial, or not-for-profit sectors.

## Ethics

Ethics approval was obtained from the South Metropolitan Health Ethics Committee (reference number: 335) Participants have given their written informed consent to participate in this study.

## Competing Interests

The authors report no competing interests.

## Notes

### Competing Interest Statement

The authors have declared no competing interest.

